# Glacier extinction homogenizes functional diversity

**DOI:** 10.1101/2023.09.25.559332

**Authors:** Nora Khelidj, Marco Caccianiga, Bruno E.L. Cerabolini, Duccio Tampucci, Gianalberto Losapio

## Abstract

**Questions:** The disappearance of glaciers threatens biodiversity and the functioning of ecosystems. To date, questions remain about the response of functional diversity to glacier extinction and its potential for adaptation to climate change. How does glacier retreat and extinction affect plant functional diversity? How do mean and variation of plant traits change with glacier retreat and extinction?

**Location:** Four glacier ecosystems in Italian Alps. Plant communities spanning 0 to ca 5,000 years on average after glacier retreat.

**Methods:** We quantify how glacier retreat affects functional diversity of plant communities analysing twelve functional traits of 117 plant species across 170 plots. First, we addressed the impact of glacier retreat on functional divergence and functional homogeneity, analysing both trait average and trait variation. Next, we explored how biodiversity (i.e., plant species richness) influenced functional diversity and how glacier retreat affected such relationship. Finally, we explored the effects of glacier retreat on mean and variation of single traits associated to carbon and nitrogen cycling and resource allocation.

**Results:** Glacier retreat homogenizes functional diversity by reducing trait variation and making communities more functionally similar. While biodiversity positively contributes to trait heterogeneity, glacier retreat erodes the support of species richness to functional diversity. We also show how glacier extinction has medium to large negative effects on the average and variation of key functional traits associated to carbon economy, but small positive effects on leaf nitrogen content.

**Conclusions:** Our results demonstrate the pervasive impact of glacier extinction on the functioning of plant communities. We stress that functional diversity and trait variation should be the focus of adaptation and mitigation actions.

## 1 INTRODUCTION

An iconic symptom of the current climate crisis is the disappearance of glaciers worldwide (Frédéric *et al*., 2015; Roe *et al*., 2017; Zemp et al., 2019; Hugonnet et al., 2021). The recent IPCC Report highlights how glaciers are unique and threatened ecological and human systems that are a major reason for concern (IPCC, 2022). Although the consequences of glacier retreat on species diversity are increasingly documented (Jacobsen *et al*., 2012; Cauvy-Fraunié & Dangles, 2019; Stibal et al., 2020; Losapio et al., 2021; Bourquin et al., 2022; Fodelianakis et al., 2022), little attention has been paid to how functional diversity would respond to glacier extinction (Caccianiga *et al*., 2006; Losapio et al., 2021). Yet, it is vitally important to understand changes in the functional diversity of these novel, fast-changing glacier environments to develop solutions for anticipating the impact of global warming on socioecological systems.

With the retreat of glaciers, new ice-free terrains are colonized by living organisms, prompting changes in species richness and composition over time (Erschbamer & Caccianiga, 2016; Ficetola et al., 2021). Previous studies indicate that species richness increases with glacier retreat in the short term (Raffl *et al*., 2006; Cauvy-Fraunié & Dangles, 2019), but this holds true only as long as glaciers are still present (Jacobsen *et al*., 2012; Stibal et al., 2020; Losapio *et al*., 2021). As glacier ecosystems comprise unique habitats that host distinctive organisms (Erschbamer & Caccianiga, 2016; Bourquin et al., 2022; Fodelianakis et al., 2022), their extinction would reduce biodiversity via both direct and indirect impacts (Losapio *et al*., 2021). Since many Alpine glaciers will disappear within the next three decades (Zekollari *et al*., 2019), we may face an up to 30 % loss in species diversity (Jacobsen *et al*., 2012; Losapio et al., 2021). However, estimates of functional diversity response remain unknown.

Approaches using functional traits can be more informative about ecological processes compared to taxonomy (Díaz & Cabido, 2001; Losapio et al., 2018; Zanzottera et al., 2020). Functional traits are any morphological, physical or phenological feature that affect the fitness of the individual (Violle *et al*., 2007). Functional traits can vary within and among plant species. One of the main cause of traits variation is the surrounding environment that filters and select traits. The combination of traits within species defines their functional type and adaptations, and reflects the abiotic and biotic environment (Díaz & Cabido, 2001). Furthermore, functional traits are also at the basis of ecosystem functioning in a given environment (Erschbamer & Caccianiga, 2016; Funk et al., 2017; Schleuning et al., 2023). Previous studies addressing the effects of glacier retreat on functional diversity reported sharp changes in trait composition related to seed dispersal (Erschbamer & Caccianiga, 2016). Following the retreat of an Alpine glacier (Rutor glacier), Caccianiga et al. (2006) reported a shift from ‘ruderal, fast growing’ species with high leaf nitrogen content to ‘stress-tolerant’ (*sensu* Grime (2001)) *plants with low N content. When considering leaf economic spectrum* traits (Reich *et al*., 2003; Wright et al., 2004), Losapio et al. (2021) documented a shift from pioneer species with ‘acquisitive’ strategies (high specific leaf area) to late species with ‘conservative’ strategies (low specific leaf area and high leaf dry matter content) along four Alpine glacier forelands. Such a turnover was accompanied by a shift from facilitative to competitive relations among plants. Similarly, Greinwald et al. (2021) reported changes in dispersal type and plant height that were consistent between two glaciers with contrasting geology. A decrease in functional divergence and homogeneity with glacier retreat suggested an increase of habitat filtering processes over niche differentiation. However, these few studies addressing the effects of glacier retreat on functional diversity have the limitations of addressing a limited number of sites or traits and, especially, considering trait average only, whereas they overlook trait variation.

Functional trait variation is often overlooked in plant community ecology, although it may be crucial for understanding key ecological processes (de Bello *et al*., 2011; Siefert et al., 2015; Losapio & Schöb, 2017; Gross et al., 2021). Functional trait variation includes the distribution or the range of traits expressed within and among plant species. The causes of trait variation range from genetic effects to environmental influence on development and phenotypic plasticity (Cornwell & Ackerly, 2009). The consequences of trait variation concern conservation genetics, ecological processes and ecosystem functioning (Cadotte *et al*., 2011; Losapio et al., 2018; Bongers et al., 2021; Carmona et al., 2021). Nevertheless, the effects of glacier retreat on functional trait variation within and between plant communities remain unknown. Although habitat protection is an important step to support biodiversity and functioning in glacier ecosystems, we also need to anticipate and predict which species to conserve or which communities to restore. To reach this goal, functional diversity can provide scalable and generalizable indications of priority species for conservation that can be tailored to the local context. Understanding functional trait variation beyond trait average is therefore key to include genetic variability, developmental constraints, and phenotypic plasticity in mitigation and adaptation strategies against glacier extinction.

Here, we addressed the following questions: (1) How does glacier extinction influence functional divergence and homogeneity, and uniqueness of functional traits – on average and considering their variation? (2) How does glacier extinction influence the relationship between biodiversity and functional diversity? (3) How does glacier extinction impact average and variation of key functional traits? In light of biodiversity decline with glacier extinction, we hypothesise that glacier extinction will be accompanied by the erosion of functional diversity. Specifically, we expect a decline in functional trait diversification, an homogenization of functional trait variation, and a loss of functional uniqueness.

## 2 METHODS

### 2.1 Field data

The study was conducted on the foreland of four glacier ecosystem sites: Vedretta d’Amola glacier, Western Trobio glacier, Rutor glacier, and Vedretta di Cedec glacier (Losapio *et al*., 2021). In each foreland, we established a transect from the glacier terminus (or glacier surface in the case of Vedretta d’Amola debris-covered glacier) to the grasslands adjacent to Little Ice Age moraines. Such transect spans terrains from recently ice-free to thousands of years old, such that plant communities range from 0 (surface of debris-covered glacier) to ca 10,000 years after glacier retreat. Terrain age was estimated as the mean between two moraines (Losapio *et al*., 2021); this way, terrains outside the LIA moraines were approximated to an average of 5,000 years old.

Hence, this transect represents a gradient of plant community development over spacetime. Furthermore, terrains outside the LIA moraines represent a scenario of glacier extinction, as opposed to glacier retreat in terrains inside the LIA moraines given the lasting impact of adjacent glacier masses. Along each transect, three to seven plots of 25–100 *m*^2^ were placed in each stage of glacier retreat (Losapio *et al*., 2021). In each plot, we surveyed plant communities by recording the presence/absence of plant species. Overall, n = 117 plant species were recorded and further analysed across four sites.

We considered plant traits relevant for plant growth, reproduction, and ecological functions (Hodgson *et al*., 1999; Díaz *et al*., 2016), namely: leaf carbon content (LCC), leaf nitrogen content (LNC), specific leaf area (SLA), leaf dry matter content (LDMC), canopy height, lateral spread, flowering start, and flowering period. We integrated our own original trait measurements with publicly available trait data (Kattge *et al*., 2020).

Traits were measured following standard protocols (Pérez-Harguindeguy *et al*., 2013). We sampled from 5 to 15 fully expanded leaves from the outer canopy of different adult healthy plants randomly selected for each species. Leaf material was stored at 4°C overnight to obtain full turgidity for determination of leaf fresh weight (LFW) and leaf area (LA). LA was determined using a digital leaf-area meter. Leaf dry weight (LDW) was then determined following drying for 24 h at 105°C. SLA was calculated as the ratio between LA and LDW. LDMC was calculated as the ratio between LDW and LFW. Leaf nitrogen concentration (LNC) and leaf carbon concentration (LCC) were quantified from dried leaf material using three randomly selected replicates that were processed with a CHNS-analyser (FlashEA 1112 series Thermo-Scientific) following the method outlined by Dalle Fratte et al. (2021). Canopy height and lateral spread were measured directly in the field. Lateral spread values were then categorised according to Hodgson et al. (1999) assigning them to one of the six categories (i.e., 1 = short-lived and non-spreading species, to 6 = widely creeping perennial species with more than 79 mm between ramets). Flowering start (FS) is defined as the month in which flowers were first produced. Flowering period (FP) is the number of months from the appearance of the first to the last flowers. Both FS and FP Hodgson et al. (1999) and were collected from Aeschimann et al. (2004).

### 2.2 Functional diversity analysis

To investigate changes in functional diversity with glacier retreat, we first considered functional divergence (FDis) and functional heterogeneity (FEve) (Mason *et al*., 2012). These two functional diversity indexes are indicators of community assembly processes and plant species coexistence (Mason *et al*., 2013). Functional divergence estimates the dispersion of the species in the multidimensional trait space, calculated as the weighted mean distance of individual species in the trait space to the centroid of all species (Laliberté & Legendre, 2010). This index is important for understanding the distribution of functions within and among plant communities. In our study context, it can shed new light on whether plant species are complementary or redundant in their functions as plant communities develop after glacier retreat. Functional heterogeneity assesses the differences between the functional traits of species in a community by measuring the degree to which plant species differ from each other in the multidimensional space of functional traits (Laliberté & Legendre, 2010). It indicates the variety of distinct ecological niches in the community and is important for understanding how plant species may interact and share resources (Schleuter *et al*., 2010) as plant communities develop. Functional diversity indices were measured with the `dbFD` function in the `FD` R package (Laliberté & Legendre, 2010).

For each trait, we considered both trait average and trait variation. Traits were averaged for each species over sites. We calculated the coefficient of variation (CV) as following: 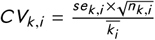, where the standard error *se* and 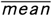 were taken for each trait *k* and for each species *i*. We calculated the CV for the following traits: LNC, LCC, CN, LA, LDW, LFW, SLA, LDMC, and CAN.

We used linear mixed-effects models to analyze the relationship between functional diversity and glacier retreat. The model was implemented using the `lmer` function in `lme4` R package (Bates *et al*., 2015). The response variables were FDIs and FEve of trait average and trait variation (four separate models), and the predictors included the quadratic effect of years of glacier retreat (log transformed), taxonomic diversity (plant species richness), the interaction between years and richness, and a random intercept for the site.

To evaluate the model output and determine the effect size, we used the `anova` function of `car` R package to perform an analysis of variance (ANOVA) on the fitted model (Fox & Weisser, 2019). We provided estimates of the model parameters and their corresponding confidence intervals using the Satterthwaite method (Kuznetsova *et al*., 2017). We also reported effect size as Cohen’s f statistic considering partial effects (Ben-Shachar *et al*., 2020). The model_parameters function

Additionally, we assessed the impact of glacier retreat on functional uniqueness of plant species. We adopted a framework based on functional dissimilarities among species for summarizing different facets of functional redundancy (Ricotta *et al*., 2016). We first calculated the pairwise trait distance matrix by means of Gower dissimilarity (`gowdis` function in (Laliberté & Legendre, 2010)). This matrix was computed for all the traits *k* (trait average only) for each site. Then, we quantified the uniqueness of species traits by addressing to what degree is a community unique as compared to a scenario where all species would be maximally dissimilar (Ricotta *et al*., 2016). We did so for each plot per site (i.e., the subset of species occurring in each plot) using the `uniqueness` function of the `adiv` R package (Pavoine, 2020). We performed a linear mixed-effects regression analysis to examine the relationship between functional uniqueness and glacier retreat. The models were built and assessed using the same procedures as described earlier.

Finally, we investigated the trait average and trait variation for each individual trait separately. For each site, we computed the observed trends in trait average and trait variation. We did so by means of the `functcomp` (Laliberté & Legendre, 2010). We performed a mixed-effects model where each trait (mean and variation) was regressed against ‘years’ using a second-degree polynomial and a random intercept for ‘site’. We reported the parameter estimates, standard errors, confidence intervals, degrees of freedom, t-values, p-values, Cohen’s f effect size, and corresponding confidence intervals.

## 3 RESULTS

### 3.1 Functional diversity indices

First, we addressed the impact of glacier retreat on functional divergence and functional homogeneity, analyzing both plant trait average and plant trait variation. We found that glacier retreat had large effects (*f* = 0.65, *p* < 0.001) on the functional divergence of trait average (Fig. 1a). These effects were negative and non-linear as functional divergence decreased exponentially with increasing glacier retreat (*r*_*l*_ = 0.22 [0.06, 0.37], *r*_*q*_ = −0.35 [-0.47, -0.19]).

**FIGURE 1.**
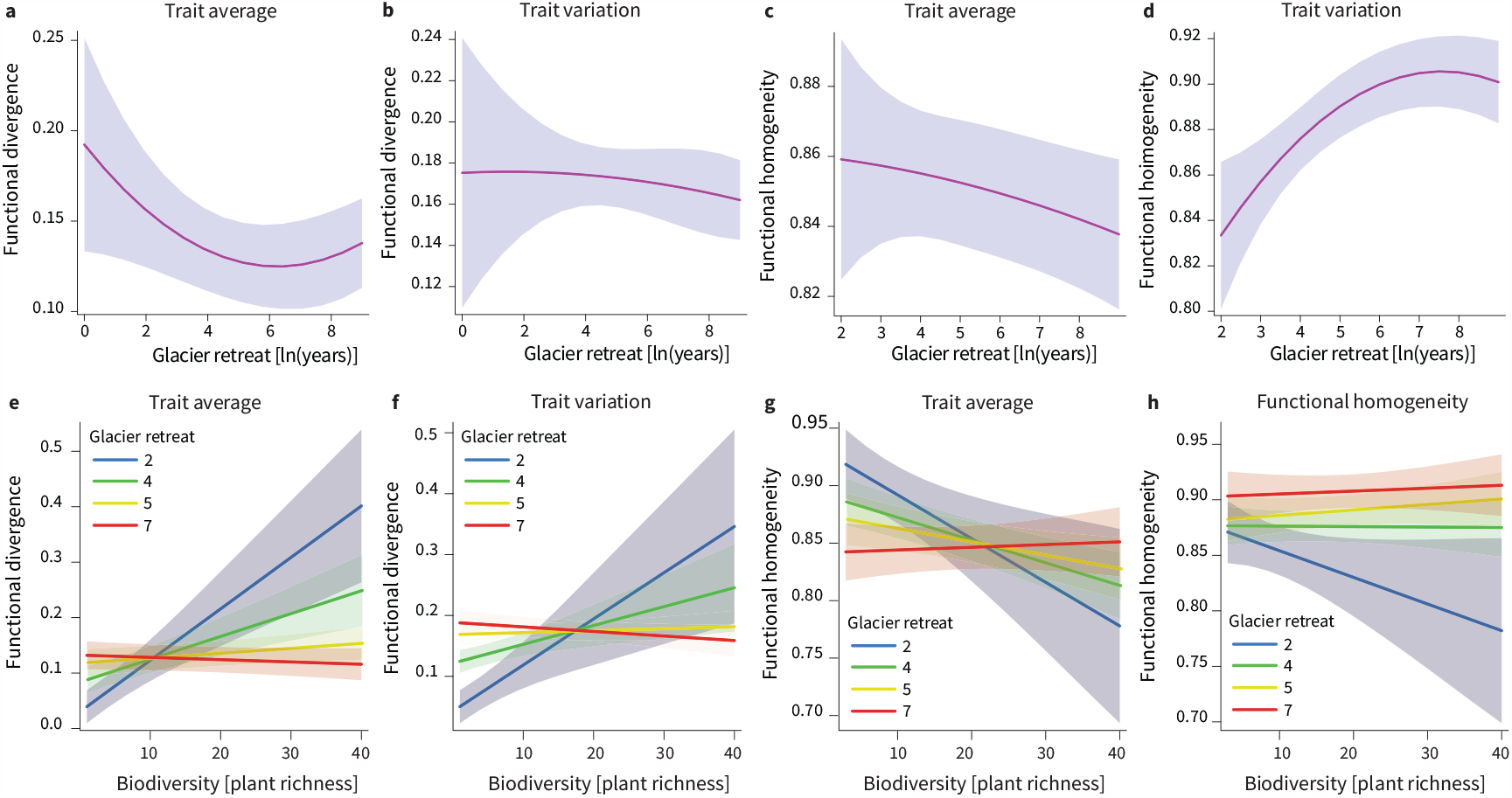
Relationship between functional diversity (*y* -axis), glacier retreat (**a:d**; log-transformed years after glacier retreat on *x* −axis), and biodiversity (**e:h**; plant species richness on *x* −axis). We report overall trends with 95% CI.

Similarly, glacier retreat had large, negative effects (*f* = 0.77, *p* < 0.001) on the functional divergence of trait variation (Fig. 1b), which also decreased exponentially with glacier retreat (*r*_*l*_ = 0.19 [0.02, 0.34], *r*_*q*_ = −0.45 [-0.56, -0.31]).

Considering the homogeneity of trait average, glacier retreat had large effects (*f* = 0.47, *p* < 0.001, Fig. 1c) that decreased homogeneity of trait variation (*r*_*l*_ = −0.42 [-0.54, -0.27], *r*_*q*_ = 0.03 [-0.14, 0.19]). Instead, glacier retreat had medium effects (*f* = 0.24, *p* < 0.025) on the functional homogeneity of trait variation (Fig. 1d). The effects of glacier retreat were negative as trait variation became more homogeneous with increasing glacier retreat (*r*_*l*_ = 0.18 [0.01, 0.34], *r*_*q*_ = 0.10 [-0.07, 0.26]).

Next, we explored how biodiversity (i.e., plant species richness) influenced functional diversity and how glacier retreat affected such relationship. We found that species richness had large effects on functional diversity (*f* = 0.27, *p* = 0.002, Extended Data Table 1) as trait average divergence increased with increasing species richness overall (*r*_*s*_ = 0.26 [0.10, 0.40]). Nevertheless, the positive effects of biodiversity on trait average divergence were eroded by glacier retreat (*r*_*l* :*s*_ = −0.28 [-0.42, -0.13], *r*_*q*:*s*_ = 0.37 [0.22, 0.49]; Fig. 1e).

Plant species richness on its own had a small effect on the functional divergence of trait variation (*f* = 0.14, *p* > 0.1, Extended Data Table 2). Yet, there was a significant interaction between glacier retreat and species richness (*f* = 0.31, *p* = 0.002) as the mainly positive relationship between biodiversity and functional divergence of trait variation (*r*_*s*_ = 0.14 [-0.04, 0.30]) was eroded by glacier retreat (*r*_*l* :*s*_ = −0.20 [-0.35, -0.04], *r*_*q*:*s*_ = 0.29 [0.13, 0.42]; Fig. 1f).

Species richness predicted homogeneity of trait average on its own (*f* = 0.26, *p* = 0.004) and depending on glacier retreat (*f* = 0.37, *p* < 0.001, Extended Data Table 3). In particular, trait variation homogeneity decreased (i.e., increase heterogeneity) with increasing species richness (*r*_*s*_ = −0.25 [-0.40, -0.08]), while glacier retreat ultimately flattened this relationship (*r*_*l* :*s*_ = 0.29 [0.12, 0.43], *r*_*q*:*s*_ = −0.03 [-0.20, 0.14]; Fig. 1g), eroding the positive role of biodiversity in community functioning. Finally, species richness had small positive effects on heterogeneity of trait variation (*r* = −0.13 [-0.31,0.07]; Extended Data Table 4) that decreased in magnitude and size with glacier retreat (*r*_*l* :*s*_ = 0.09 [-0.08, 0.25], *r*_*q*:*s*_ = −0.20 [-0.35, -0.03]; Fig. 1h).

Finally, we considered how functional uniqueness changed with climate change (Extended Data Table 5). We found that glacier retreat had strong statistical effects on functional uniqueness (*f* = 1.132, *p* < 0.001) as the functional uniqueness of plant species decreased with increasing glacier retreat (*r*_*l*_ = −0.46 [-0.57, -0.32], *r*_*q*_ = 0.57 [0.46, 0.66]; Fig. 2). Furthermore, functional uniqueness decreased with increasing species richness (*r*_*s*_ = −0.50 [-0.61, -0.36]), an effect also mediated by glacier retreat (*r*_*l* :*s*_ = 0.43 [0.29, 0.5], *r*_*q*:*s*_ = −0.45 [-0.56, -0.31]; Fig. 2).

**FIGURE 2.**
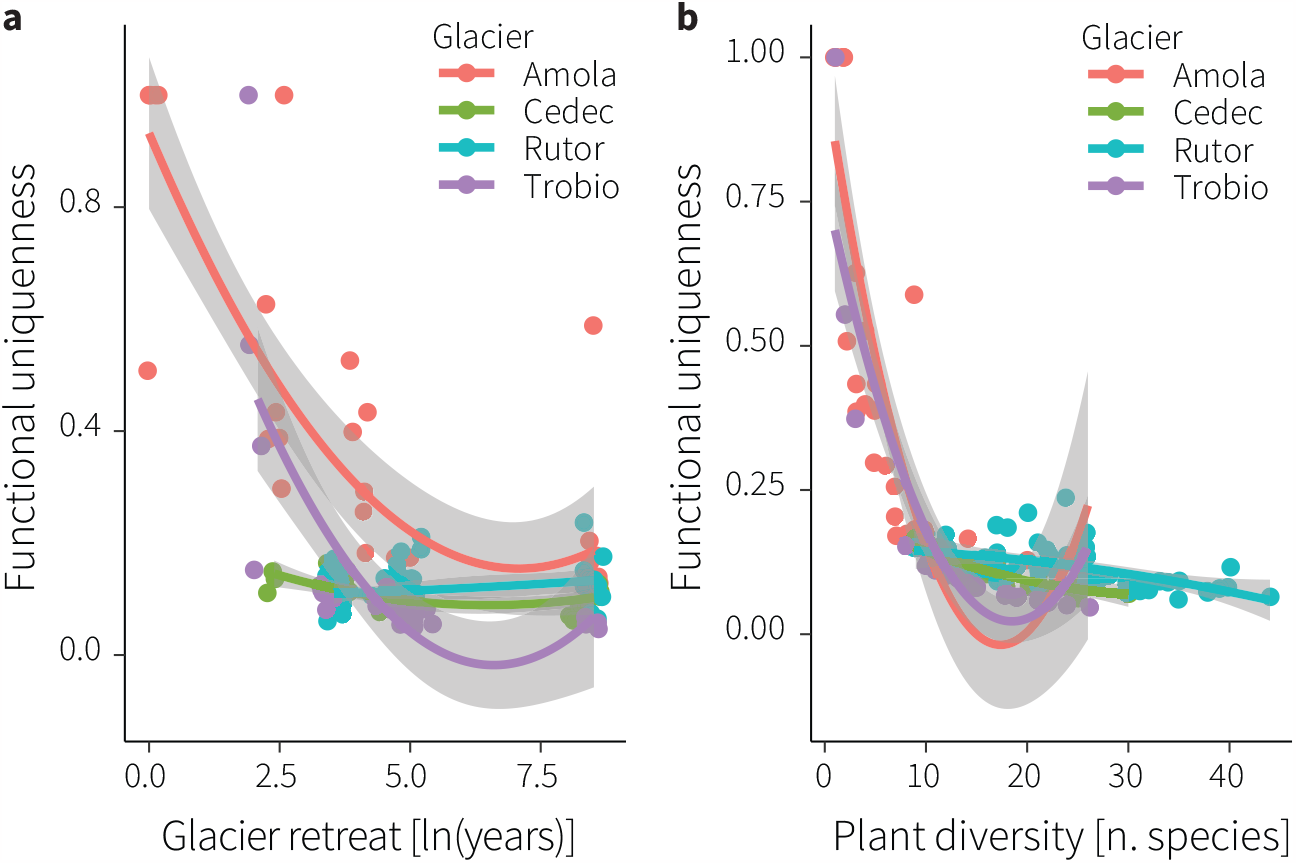
Relationship between functional uniquenness (*y* -axis), glacier retreat (**a**; log-transformed years after glacier retreat on *x* −axis), and biodiversity (**b**; plant species richness on *x* −axis). We report trends with 95% CI among sites.

### 3.2 Single traits

We proceeded with examining how single plant traits changed with glacier retreat (Fig. 4; Extended Data Table 6 and Extended Data Table 7). Leaf carbon content average increased with glacier retreat (*r*_*l*_ = 0.726 [0.649, 0.783], *r*_*q*_ = 0.334 [0.183, 0.462]). On the contrary, leaf carbon content variation tended to decrease with glacier extinction (*r*_*l*_ = 0.111 [-0.052, 0.266], *r*_*q*_ = −0.418 [-0.533, -0.276]). Leaf nitrogen content average increased initially but ultimately decreased with glacier extinction (*r*_*l*_ = 0.047 [-0.115, 0.206], *r*_*q*_ = −0.338 [-0.465, -0.188]), whereas leaf nitrogen content variation increased with glacier extinction (*r*_*l*_ = 0.177 [0.017, 0.324]). The average carbon to nitrogen content ratio showed a positive parabolic change with glacier retreat (*r*_*l*_ = 0.221 [0.060, 0.364], *r*_*q*_ = −0.372 [0.226, 0.494]), while the variation in carbon to nitrogen content ratio showed a negative parabolic change with glacier retreat (*r*_*l*_ = 0.290 [0.134, 0.424], *r*_*q*_ = −0.391 [-0.512, -0.246]).

**FIGURE 3.**
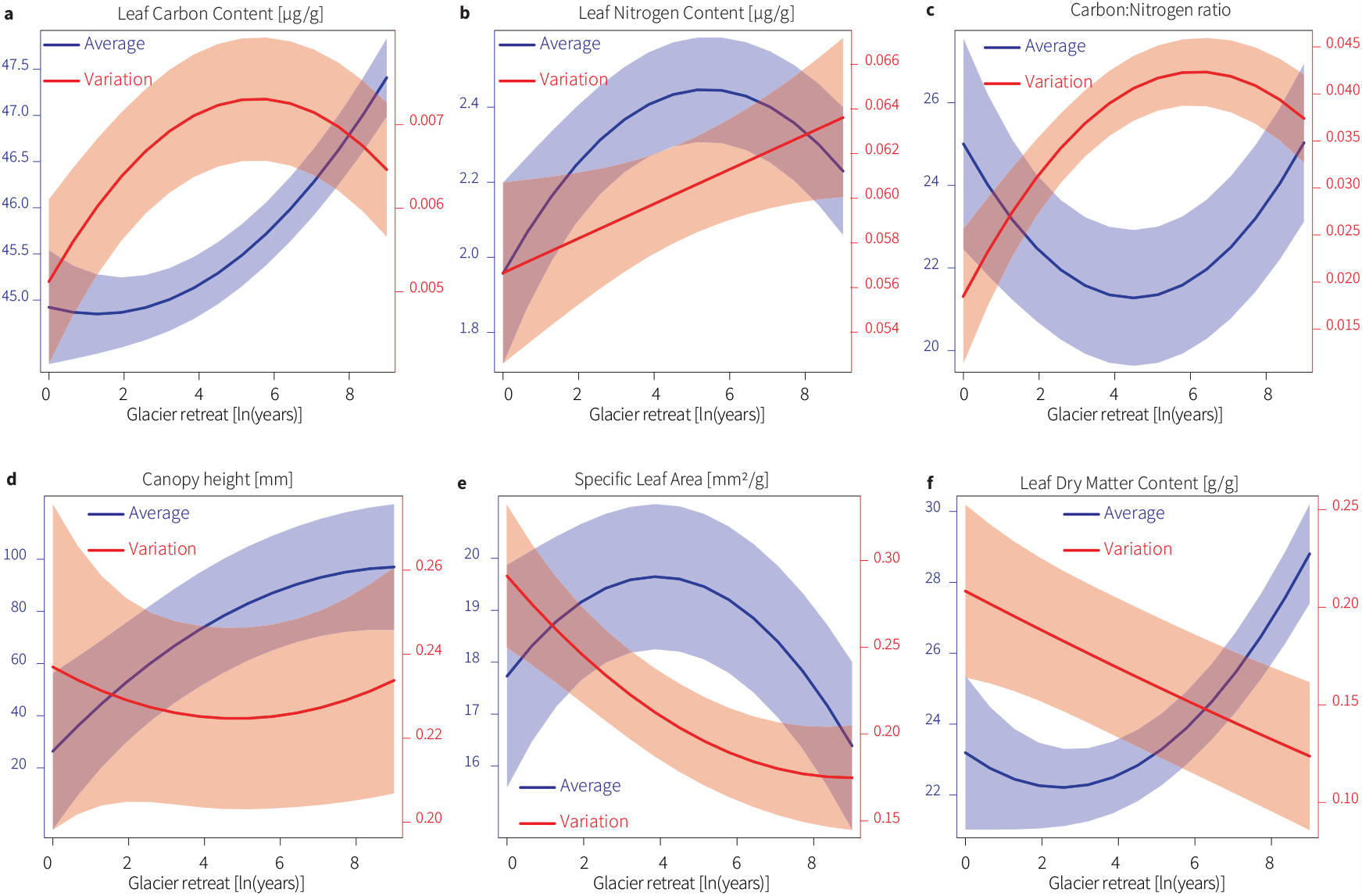
Relationship between plant traits (trait average in blue; trait variation in orange; *y* -axis) and glacier retreat (log-transformed years after glacier retreat, *x* −axis). We report overall trends with 95% CI.

**FIGURE 4.**
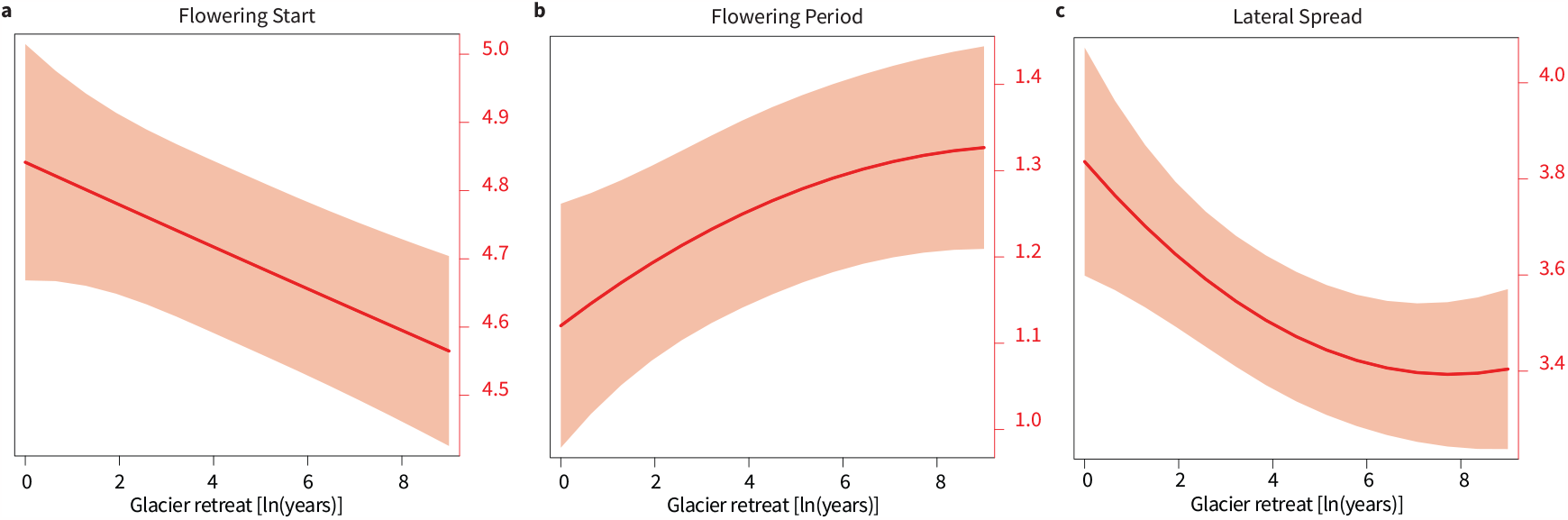
Relationship between flowering strategy (**a**; *y* -axis), flowering period (**b**; *y* -axis), lateral growth (**c**; *y* -axis), and glacier retreat (log-transformed years after glacier retreat, *x* −axis). We report overall trends with 95% CI.

**FIGURE 5.**
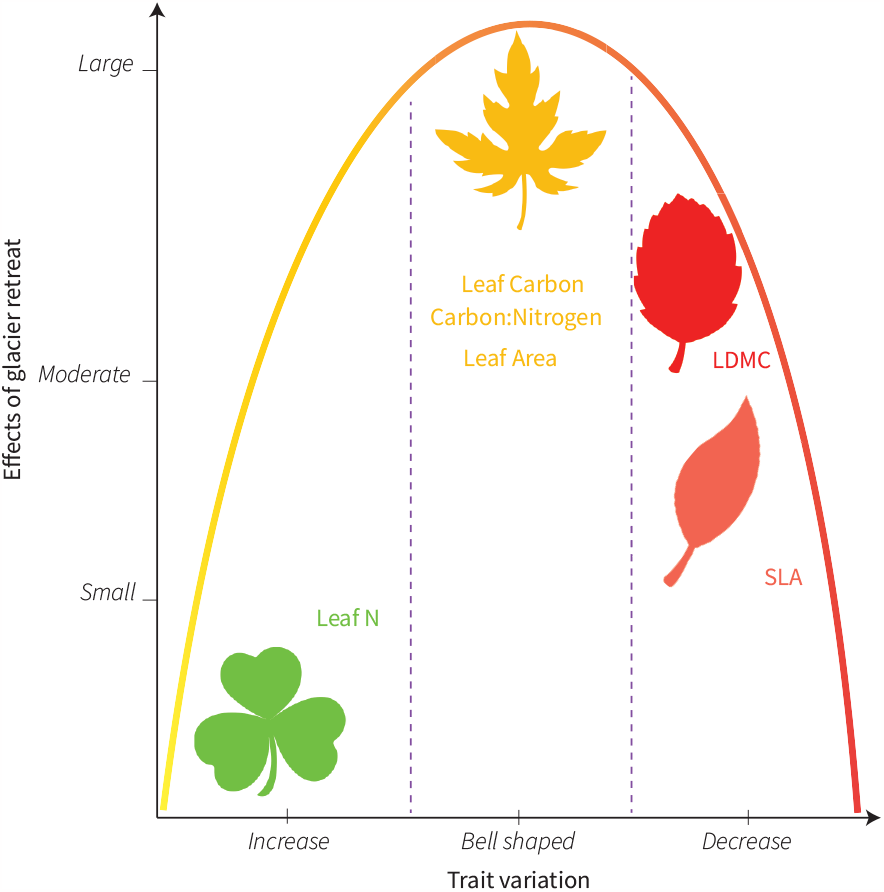
Summary of glacier retreat effects (direction and strength) on the variation of plant functional traits.

Considering ‘leaf economic spectrum’ traits, SLA average showed a negative parabolic change with glacier retreat (*r*_*l*_ = −0.314 [-0.446, -0.160], *r*_*q*_ = −0.310 [-0.442, -0.157]) while SLA variation decreased exponentially with glacier retreat (*r*_*l*_ = −0.474 [-0.580, -0.341], *r*_*q*_ = 0.186 [0.025, 0.332]). On the contrary, LDMC average increased with glacier retreat (*r*_*l*_ = 0.577 [0.463, 0.663], *r*_*q*_ = 0.329 [0.176, 0.459]), whereas LDMC variation sharply decreased (*r*_*l*_ = 0. − 497 [-0.599, -0.367]).

Notably, average canopy height tended to increase with glacier retreat (*r*_*l*_ = 0.464 [0.329, 0.572], *r*_*q*_ = −0.166 [-0.315, -0.004]). On the contrary, lateral spread decreased with glacier retreat (*r*_*l*_ = −0.294 [-0.429, -0.139], *r*_*q*_ = −0.157 [-0.005, 0.306]; Fig. 4).

Finally, we also found that flowering start sharply decreased with glacier extinction (*r*_*l*_ = −0.386 [-0.507, -0.241]; Fig. 4a), while flowering period increased with glacier retreat (*r*_*l*_ = 0.336 [0.185, 0.465]; Fig. 4b; Extended Data Table 6).

## 4 DISCUSSION

Our results indicate that the retreat of glaciers is reducing the functional diversification of plant communities. By making communities more similar to each other in terms of average functions and functional variation, glacier extinction reduces the redundancy of functions. They also indicate that glacier extinctions will make functional variation among species more homogeneous, with potential implications for local adaptation to climate change.

These results indicate that the extinction of glaciers negatively impacts functional diversity by decreasing both niche differentiation and the variation in ecological functions performed by diverse, species-rich communities. Taken together, glacier retreat had dual impacts on functional diversity: directly, it decreases and homogenizes functional diversification, and, indirectly, it erodes the positive effects of species richness on functional divergence and heterogeneity. Consistent with results of models and recent empirical work (Cadotte *et al*., 2011; Griffin-Nolan et al., 2019; Song *et al*., 2014), ecosystem functions and possibilities for local adaptation may be accordingly reduced too. When focusing on single traits, our results further demonstrate that glacier retreat strongly impacts trait average and trait variation. Trait average for each single trait changes with glacier retreat either linearly or non-linearly. Instead, trait variation tends to decrease for most of the analyzed traits.

### 4.1 Community processes and diversity–functioning relationship

To understand the influence of glacier retreat on community structure and community processes, we used different trait-based metrics, namely functional divergence and functional evenness, considering both trait average and trait variation. Most productive ecosystems are hypothesized to host more diverse species and mostly divergent traits (Song et al., 2014). This is mainly attributed to a decrease in niche overlap due to the high diversity of function (Cadotte *et al*., 2011; Song et al., 2014) and an increase in competition, which prevents species posing too similar traits to co-exist while maintaining high level of taxonomic and functional diversity (Greinwald *et al*., 2021; Grime, 2006). Species-rich communities are mainly characterized by a high functional evenness, as most of the niche space is filled (Mouchet *et al*., 2010), and high functional divergence, which reflects the uniqueness trait and diversity of traits occurring in the habitat (Mouchet *et al*., 2010).

Our results indicate that functional divergence and evenness of trait average are the highest in early stages and decrease with glacier retreat. On the contrary, we observed lower values of functional evenness and divergence at later stages in old communities. Functional divergence relates to the abundance of traits. High values of divergence are observed when the most abundant traits show very different values, which suggests a high variability in trait distribution. Low divergence values occur when the abundant trait shows similar values. However, habitat filtering is usually observed in harsh habitats where specific sets of traits are needed for species to survive (Greinwald *et al*., 2021; Mouchet *et al*., 2010). Early colonisation stages or young moraines are known to host harsh environments, where climatic conditions are less suitable or nutrient-poor soil (Hågvar *et al*., 2020; Khedim et al., 2021). Late colonisation stage or older moraines are more productive with, high biodiversity, nutrients rich soil, and often climatic condition less extreme than toward younger moraines (Greinwald *et al*., 2021; Losapio et al., 2021). Following that, we expect to observe that the main process driving community structure is habitat filtering in the early stages, then shifting to competitive exclusion in later stages. Yet, our results indicate a contradictory pattern to what can be expected from community assembly theory concerning colonisation processes in glacier forelands.

We found that functional divergence was relatively low, suggesting that filtering might be happening all along the foreland, with varying degree depending on terrain age. Greinwald et al. (2021) have done a similar study and found similar results, suggesting that the alpine climate itself is harsher than lowland climate. This climate already acts as strong limiting factors for plant establishment and survival. Yet, we found that the filtering process is stronger in the latest stages, suggesting that the environment is less suitable for plants to inhabit. We also observed a sharp increase in plant height in our results. Tussock-dominated patches and shrublands can act as environmental filters, by altering environmental conditions, mainly by modifying light availability. With the increase in plant height, only a limited number of plants have access to maximum sunlight. Light thus becomes a limiting factor for the growth and survival of plant species (Depauw *et al*., 2020; Happonen et al., 2021). These results suggest that light limitation exerted by shrubs (e.g. *Rhododendron ferrugineum*) and tussock grasses/sedges (e.g. *Carex curvula*) may be a potential mechanism underlying the exclusion of herbaceous, pioneer species and driving biodiversity decline with glacier extinction.

We further decomposed the effects of species richness and glacier retreat on functional diversity. It is generally assumed that the highest the biodiversity, the highest the functional diversity, and thus the divergence between traits (Cadotte *et al*., 2011). Here, the hypothesis holds true in the early stages of community assembly. We found that with increasing time, the influence of biodiversity on traits is eroded by glacier retreat as the positive relationship between species richness and functional traits is not observed anymore. This flatten diversity–functioning relationship suggests that functional diversity and trait similarity change no matter the number of species present in the community. Furthermore, we also observed a reduction of unique traits with glacier retreat. A possible explanation is that the rate of decrease in functional diversity is much stronger that the taxonomic diversity decrease rate. As environmental filtering may allow different species with similar traits to coexist, glacier retreat would lead to a faster decrease in functional diversity than taxonomic diversity.

### 4.2 Single traits

We further assess the variation of single traits after glacier retreat. Both leaf carbon content and leaf nitrogen content show dramatic variation along the foreland. With glacier retreat, leaf carbon and nitrogen content increase at first. But ultimately nitrogen content decreases in the oldest communities, while carbon doesn’t. This divergence results in the initial decrease of C:N ratio after deglaciation but in an ultimate increase in the oldest communities. These results are similar to trends observed in soil nutrients too (Khedim *et al*., 2021). Leaf nutrient concentration reflects the nutrient concentration in the surrounding environment (Zhang *et al*., 2020). Early stages of deglaciation are characterized by parent material and nutrient-poor soil (Khedim *et al*., 2021). Accordingly, our results show that leaves have a high C:N ratio in pioneer communities. The process is comparable to the early stage of terrestrial plant evolution when soil presented similar characteristics. Plants evolved a high C:N ratio to be able to survive in strongly poor nutrient environments and resist environmental stress such as cold environments(Zhang *et al*., 2020). With time, soil becomes richer due to weathering from minerals, the decomposition of plants and the increase of organisms able to fix atmospheric N (Khedim *et al*., 2021; Zhang et al., 2020). A similar process happens after the retreat of glaciers and it is highlighted by our results of C:N ratio decreasing with time. Our results are consistent with previous ones (Zhang *et al*., 2020)demonstrating that plant C:N ratio decreases due to a nutrient-rich soil increasing leaves concentration of N and a decrease of leaves C due to high competition for light.

This trend is further corroborated by the analyses of traits of the leaf economic spectrum, primarily SLA and LDMC(Wright *et al*., 2004). We found that plants with high SLA and low LDMC increase during early community assembly stages, indicating that plants with fast growth strategy are associated with early colonization. Colonizers are replaced toward the latest stages by species with low SLA and high LDMC (denser leaves and slow growers), typically more stress-tolerant species(Erschbamer & Caccianiga, 2016; Losapio et al., 2021). Stress-tolerant species can persist in time by outcompeting pioneer species (Losapio *et al*., 2021). Taken together, these results suggest that different forces are driving the community structure of plants and trait distribution of plant communities in the aftermath of glacier retreat.

### 4.3 The importance of trait diversity and variation

Our study highlighted the negative impact of glacier retreat on functional diversity, even when biodiversity increased. It is assumed that species diversity and functional diversity are tightly linked such that when one increases so does the other (Cadotte *et al*., 2011). However, the effect of glacier retreat is becoming more important with time eroding the positive effect of species diversity on functional diversity. It is therefore important to consider functional diversity when assessing community change, not only in the context of glacier retreat but also in other communities that are changing and reassembling fast and in novel, unexpected ways.

Functional divergence tends to decrease in stressful environments. For many years, plants and living organisms have evolved and adapted suitable traits to survive in their changing environment. However, with the recent retreat of glaciers at unprcedented rates (Zemp *et al*., 2015), biotic and abiotic conditions change rapidly in glacier foreland, posing serious challenges to evolution and constrains to adapation. Plants are not necessarily able to adapt fast enough to the new environmental and climatic conditions. Not only plants do not adapt quickly enough to those new environments, but the decrease in trait divergence and variation further leads to a decrease in their ability to adapt to future climates and environments (Rodman *et al*., 2021). This process is creating a vicious circle exposing more and more species to extinction risk. There are thus increasing studies showing the necessity to study and promote functional traits (Roches *et al*., 2017; Heilmeier, 2019). Our results highlight the importance of accounting for functional divergence and functional variation in conservation strategies aimed at anticipating the impact of glacier retreat on biodiversity.

Furthermore, our study also shows that the variation of most individual traits also decreases with glacier retreat, with the exception of leaf nitrogen and plant height. There is consensus on the importance of maintaining a high level of functional divergence, diversity, and redundancy to promote ecosystem processes and support resilience (Díaz & Cabido, 2001; Tilman, 1997; Ricotta et al., 2016; Westerband et al., 2021). Westerband et al. (2021) have previously demonstrated that mixed stands (in that case associated with a higher functional diversity) have better resilience and resistance to drought events in forest ecosystems. The underlying process is mainly due to a better resource repartitions and lower niche overlap among species. An example of better niche partitioning thanks to functional traits is canopy size as a mixture of different canopy sizes enables trees to maximize light intake (Forrester *et al*., 2018). In addition, Chen & Chen (2021) have demonstrated that increasing plant functional diversity is helping the balance in soil nutrients and nutrient cycling. Traits are a result of genetic variation and phenotypic plasticity reflecting the surrounding environment. Part of the diversity is thus inheritable (Siefert *et al*., 2015). There are increasing studies showing the necessity of protecting and enhancing genetic diversity (Hoban *et al*., 2020; Teixeira & Huber, 2021). Likewise different alleles of the same genes are necessary in natural environments, also different versions of the same traits are also necessary (Roches *et al*., 2017; Heilmeier, 2019), stressing the need of addressing functional variation besides trait average.

In conclusion, different forces drive plant community assembly after glacier retreat. The majority of plant functional traits are altered by the rapid retreat of glaciers and climate change more than by changes in species richness. Measures are necessary to maintain a high level of functional diversity, or we face a decline in many alpine, cold-adapted species. Functional traits are both inherited and driven by the environment. It is thus necessary to consider both genetic diversity, habitat structure and community composition when protecting and supporting functional diversity.

## 5 ACKNOWLEDGMENTS

We thank Chiara Maffioletti for her help during fieldwork and the Biodiversity Change group for fruitful discussions.

## 6 AUTHOR CONTRIBUTIONS

GL designed the research. GL, MC, BELC, and DT collected data. NK and GL analyzed the data. NK and GL wrote the manuscript. All authors contributed to manuscript editing, read and approved the final manuscript.https://www.overleaf.com/project/

## 7 DATA AVAILABILITY

The data collected for this study are deposited on GitHub https://github.com/losapio/JVS-Glacier-extinction-homogenizes-plant-functional-diversity.git. The R script (no novel code) to reproduce the analyses and figures is also provided on GitHub and as electronic Supplementary Information.

## 8 ETHICS DECLARATIONS

The authors declare no competing interests.

## 9 SUPPORTING INFORMATION

The R code with supplementary figures and tables can be found in the online version of the paper.

## References

Aeschimann, D., Lauber, K., Moser, D. & Theurillat, J.P. (2004) Flora alpina. Haupt, Bern.

Bates, D., Mächler, M., Bolker, B. & Steve, W. (2015) Fitting Linear Mixed-Effects Models using lme4. Journal of Statistical Software 67, 1–48.

Ben-Shachar, M., Lüdecke, D. & D M. (2020) effectsize: Estimation of effect size indices and standardized parameter. Journal of Open Source Software 5, 2815.

Bongers, F.J., Schmid, B., Bruelheide, H., Bongers, F., Li, S., von Oheimb, G., Li, Y., Cheng, A., Ma, K. & Liu, X. (2021) Functional diversity effects on productivity increase with age in a forest biodiversity experiment. Nature Ecology & Evolution 5, 1594– 1603.

Bourquin, M., Busi, S.B., Fodelianakis, S., Peter, H., Washburne, A., Kohler, T.J., Ezzat, L., Michoud, G., Wilmes, P. & Battin, T.J. (2022) The microbiome of cryospheric ecosystems. Nature Communications 13, 3087.

Caccianiga, M., Luzzaro, A., Pierce, S., Ceriani, R.M. & Cerabolini, B. (2006) The functional basis of a primary succession resolved by CSR classification. Oikos 112, 10–20.

Cadotte, M.W., Carscadden, K. & Mirotchnick, N. (2011) Beyond species: Functional diversity and the maintenance of ecological processes and services. Journal of Applied Ecology 48, 1079–1087.

Carmona, C.P., Tamme, R., Pärtel, M., de Bello, F., Brosse, S., Capdevila, P., González-M, R., González-Suárez, M., Salguero-Gómez, R., Vásquez-Valderrama, M. & Toussaint, A. (2021) Erosion of global functional diversity across the tree of life. Science Advances 7, eabf2675.

Cauvy-Fraunié, S. & Dangles, O. (2019) A global synthesis of biodiversity responses to glacier retreat. Nature Ecology & Evolution 3, 1675–1685.

Cornwell, W.K. & Ackerly, D.D. (2009) Community assembly and shifts in plant trait distributions across an environmental gradient in coastal California. Ecological Monographs 79, 109–126.

Dalle Fratte, M., Pierce, S., Zanzottera, M. & Cerabolini, B.E.L. (2021) The association of leaf sulfur content with the leaf economics spectrum and plant adaptive strategies. Functional Plant Biology 48, 924–935.

de Bello, F., Lavorel, S., Albert, C.H., Thuiller, W., Grigulis, K., Dolezal, J., Janeček, Š. & Lepš, J. (2011) Quantifying the relevance of intraspecific trait variability for functional diversity. Methods in Ecology and Evolution 2, 163–174.

Depauw, L., Perring, M.P., Landuyt, D., Maes, S.L., Blondeel, H., Lombaerde, E.D., Brūmelis, G., Brunet, J., Closset-Kopp, D., Czerepko, J., Decocq, G., den Ouden, J., Gawryś, R., Härdtle, W., Hédl, R., Heinken, T., Heinrichs, S., Jaroszewicz, B., Kopecký, M., Liepina, I., Macek, M., Máliš, F., Schmidt, W., Smart, S.M., Ujházy, K., Wulf, M. & Verheyen, K. (2020) Light availability and land-use history drive biodiversity and functional changes in forest herb layer communities. Journal of Ecology 108, 1411–1425.

Díaz, S., Kattge, J., Cornelissen, J.H.C., Wright, I.J., Lavorel, S., Dray, S., Reu, B., Kleyer, M., Wirth, C., Colin Prentice, I., Garnier, E., Bönisch, G., Westoby, M., Poorter, H., Reich, P.B., Moles, A.T., Dickie, J., Gillison, A.N., Zanne, A.E., Chave, J., Joseph Wright, S., Sheremet’ev, S.N., Jactel, H., Baraloto, C., Cerabolini, B., Pierce, S., Shipley, B., Kirkup, D., Casanoves, F., Joswig, J.S., Günther, A., Falczuk, V., Rüger, N., Mahecha, M.D. & Gorné, L.D. (2016) The global spectrum of plant form and function. Nature 529, 167–171.

Díaz, S. & Cabido, M. (2001) Vive la différence: plant functional diversity matters to ecosystem processes. Trends in Ecology Evolution 16, 646–655.

Erschbamer, B. & Caccianiga, M.S. (2016) Glacier Forelands: Lessons of Plant Population and Community Development, vol. Progress in Botany 78, pp. 259–284. Springer International Publishing, Cham.

Ficetola, G.F., Marta, S., Guerrieri, A., Gobbi, M., Ambrosini, R., Fontaneto, D., Zerboni, A., Poulenard, J., Caccianiga, M. & Thuiller, W. (2021) Dynamics of ecological communities following current retreat of glaciers. Annual Review of Ecology, Evolution, and Systematics 52, 405–426.

Fodelianakis, S., Washburne, A.D., Bourquin, M., Pramateftaki, P., Kohler, T.J., Styllas, M., Tolosano, M., De Staercke, V., Schön, M., Busi, S.B., Brandani, J., Wilmes, P., Peter, H. & Battin, T.J. (2022) Microdiversity characterizes prevalent phylogenetic clades in the glacier-fed stream microbiome. The ISME Journal 16, 666–675.

Forrester, D.I., Ammer, C., Annighöfer, P.J., Barbeito, I., Bielak, K., Bravo-Oviedo, A., Coll, L., del Río, M., Drössler, L., Heym, M., Hurt, V., Löf, M., den Ouden, J., Pach, M., Pereira, M.G., Plaga, B.N.E., Ponette, Q., Skrzyszewski, J., Sterba, H., Svoboda, M., Zlatanov, T.M. & Pretzsch, H. (2018) Effects of crown architecture and stand structure on light absorption in mixed and monospecific <i>fagus sylvatica</i> and <i>pinus sylvestris</i> forests along a productivity and climate gradient through europe. Journal of Ecology 106, 746–760.

Fox, J. & Weisser, S. (2019) An R companion to Applied Regression. Sage, Thousand Oaks, CA, 3rd edn.

Frédéric, H., Olivier, B., Mattia, B. N. L.S., Sébastien, L., Thierry, A. Y. L.J.Y., Jean-Philippe, A. & C. C.S. (2015) Erosion by an alpine glacier. Science 350, 193–195.

Funk, J.L., Larson, J.E., Ames, G.M., Butterfield, B.J., Cavender-Bares, J., Firn, J., Laughlin, D.C., Sutton-Grier, A.E., Williams, L. & Wright, J. (2017) Revisiting the <scp>h</scp> oly <scp>g</scp> rail: using plant functional traits to understand ecological processes. Biological Reviews 92, 1156–1173.

Greinwald, K., Gebauer, T., Musso, A. & Scherer-Lorenzen, M. (2021) Similar successional development of functional community structure in glacier forelands despite contrasting bedrocks. Journal of Vegetation Science 32, e12993.

Griffin-Nolan, R.J., Blumenthal, D.M., Collins, S.L., Farkas, T.E., Hoffman, A.M., Mueller, K.E., Ocheltree, T.W., Smith, M.D., Whitney, K.D. & Knapp, A.K. (2019) Shifts in plant functional composition following long-term drought in grasslands. Journal of Ecology 107, 2133–2148.

Grime, J.P. (2001) Plant strategies, vegetation processes, and ecosystem properties. Wiley, 2nd edn.

Grime, J.P. (2006) Trait convergence and trait divergence in herbaceous plant communities: Mechanisms and consequences. Journal of Vegetation Science 17, 255–260.

Gross, N., Le Bagousse-Pinguet, Y., Liancourt, P., Saiz, H., Violle, C. & Munoz, F. (2021) Unveiling ecological assembly rules from commonalities in trait distributions. Ecology Letters 24, 1668–1680.

Happonen, K., Muurinen, L., Virtanen, R., Kaakinen, E., Grytnes, J., Kaarlejärvi, E., Parisot, P., Wolff, M. & Maliniemi, T. (2021) Trait-based responses to land use and canopy dynamics modify long-term diversity changes in forest understories. Global Ecology and Biogeography 30, 1863–1875.

Heilmeier, H. (2019) Functional traits explaining plant responses to past and future climate changes. Flora 254, 1–11.

Hoban, S., Bruford, M., Jackson, J.D., Lopes-Fernandes, M., Heuertz, M., Hohenlohe, P.A., Paz-Vinas, I., Sjögren-Gulve, P., Segelbacher, G., Vernesi, C., Aitken, S., Bertola, L.D., Bloomer, P., Breed, M., Rodríguez-Correa, H., Funk, W.C., Grueber, C.E., Hunter, M.E., Jaffe, R., Liggins, L., Mergeay, J., Moharrek, F., O’Brien, D., Ogden, R., Palma-Silva, C., Pierson, J., Ramakrishnan, U., Simo-Droissart, M., Tani, N., Waits, L. & Laikre, L. (2020) Genetic diversity targets and indicators in the cbd post-2020 global biodiversity framework must be improved. Biological Conservation 248, 108654.

Hodgson, J., Wilson, P., Hunt, R., Grime, J. & Thompson, K. (1999) Allocating C-S-R plant functional types: a soft approach to a hard problem. Oikos 85, 282–294.

Hugonnet, R., McNabb, R., Berthier, E., Menounos, B., Nuth, C., Girod, L., Farinotti, D., Huss, M., Dussaillant, I., Brun, F. & Kääb, A. (2021) Accelerated global glacier mass loss in the early twenty-first century. Nature 592, 726–731.

Hågvar, S., Gobbi, M., Kaufmann, R., Ingimarsdóttir, M., Caccianiga, M., Valle, B., Pantini, P., Fanciulli, P.P. & Vater, A. (2020) Ecosystem birth near melting glaciers: A review on the pioneer role of ground-dwelling arthropods. Insects 11, 644.

IPCC (2022) Climate Change 2022: Impacts, Adaptation, and Vulnerability., vol. Contribution of Working Group II to the Sixth Assessment Report of the Intergovernmental Panel on Climate Change. Cambridge University Press.

Jacobsen, D., Milner, A.M., Brown, L.E. & Dangles, O. (2012) Biodiversity under threat in glacier-fed river systems. Nature Climate Change 2, 361–364.

Kattge, J., Bönisch, G., Díaz, S., Lavorel, S., Prentice, I.C., Leadley, P., Tautenhahn, S., Werner, G.D.A., Aakala, T., Abedi, M., Acosta, A.T.R., Adamidis, G.C., Adamson, K., Aiba, M., Albert, C.H., Alcántara, J.M., Alcázar C, C., Aleixo, I., Ali, H., Amiaud, B., Ammer, C., Amoroso, M.M., Anand, M., Anderson, C., Anten, N., Antos, J., Apgaua, D.M.G., Ashman, T.L., Asmara, D.H., Asner, G.P., Aspinwall, M., Atkin, O., Aubin, I., Baastrup-Spohr, L., Bahalkeh, K., Bahn, M., Baker, T., Baker, W.J., Bakker, J.P., Baldocchi, D., Baltzer, J., Banerjee, A., Baranger, A., Barlow, J., Barneche, D.R., Baruch, Z., Bastianelli, D., Battles, J., Bauerle, W., Bauters, M., Bazzato, E., Beckmann, M., Beeckman, H., Beierkuhnlein, C., Bekker, R., Belfry, G., Belluau, M., Beloiu, M., Benavides, R., Benomar, L., Berdugo-Lattke, M.L., Berenguer, E., Bergamin, R., Bergmann, J., Bergmann Carlucci, M., Berner, L., Bernhardt-Römermann, M., Bigler, C., Bjorkman, A.D., Blackman, C., Blanco, C., Blonder, B., Blumenthal, D., Bocanegra-González, K.T., Boeckx, P., Bohlman, S., Böhning-Gaese, K., Boisvert-Marsh, L., Bond, W., Bond-Lamberty, B., Boom, A., Boonman, C.C.F., Bordin, K., Boughton, E.H., Boukili, V., Bowman, D.M.J.S., Bravo, S., Brendel, M.R., Broadley, M.R., Brown, K.A., Bruelheide, H., Brumnich, F., Bruun, H.H., Bruy, D., Buchanan, S.W., Bucher, S.F., Buchmann, N., Buitenwerf, R., Bunker, D.E., Bürger, J., Burrascano, S., Burslem, D.F.R.P., Butterfield, B.J., Byun, C., Marques, M., Scalon, M.C., Caccianiga, M., Cadotte, M., Cailleret, M., Camac, J., Camarero, J.J., Campany, C., Campetella, G., Campos, J.A., Cano-Arboleda, L., Canullo, R., Carbognani, M., Carvalho, F., Casanoves, F., Castagneyrol, B., Catford, J.A., Cavender-Bares, J., Cerabolini, B.E.L., Cervellini, M., Chacón-Madrigal, E., Chapin, K., Chapin, F.S., Chelli, S., Chen, S.C., Chen, A., Cherubini, P., Chianucci, F., Choat, B., Chung, K.S., Chytrý, M., Ciccarelli, D., Coll, L., Collins, C.G., Conti, L., Coomes, D., Cornelissen, J.H.C., Cornwell, W.K., Corona, P., Coyea, M., Craine, J., Craven, D., Cromsigt, J.P.G.M., Csecserits, A., Cufar, K., Cuntz, M., da Silva, A.C., Dahlin, K.M., Dainese, M., Dalke, I., Dalle Fratte, M., Dang-Le, A.T., Danihelka, J., Dannoura, M., Dawson, S., de Beer, A.J., De Frutos, A., De Long, J.R., Dechant, B., Delagrange, S., Delpierre, N., Derroire, G., Dias, A.S., Diaz-Toribio, M.H., Dimitrakopoulos, P.G., Dobrowolski, M., Doktor, D., Dřevojan, P., Dong, N., Dransfield, J., Dressler, S., Duarte, L., Ducouret, E., Dullinger, S., Durka, W., Duursma, R., Dymova, O., E-Vojtkó, A., Eckstein, R.L., Ejtehadi, H., Elser, J., Emilio, T., Engemann, K., Erfanian, M.B., Erfmeier, A., Esquivel-Muelbert, A., Esser, G., Estiarte, M., Domingues, T.F., Fagan, W.F., Fagúndez, J., Falster, D.S., Fan, Y., Fang, J., Farris, E., Fazlioglu, F., Feng, Y., Fernandez-Mendez, F., Ferrara, C., Ferreira, J., Fidelis, A., Finegan, B., Firn, J., Flowers, T.J., Flynn, D.F.B., Fontana, V., Forey, E., Forgiarini, C., François, L., Frangipani, M., Frank, D., Frenette-Dussault, C., Freschet, G.T., Fry, E.L., Fyllas, N.M., Mazzochini, G.G., Gachet, S., Gallagher, R., Ganade, G., Ganga, F., García-Palacios, P., Gargaglione, V., Garnier, E., Garrido, J.L., de Gasper, A., Gea-Izquierdo, G., Gibson, D., Gillison, A.N., Giroldo, A., Glasenhardt, M.C., Gleason, S., Gliesch, M., Goldberg, E., Göldel, B., Gonzalez-Akre, E., Gonzalez-Andujar, J.L., González-Melo, A., González-Robles, A., Graae, B.J., Granda, E., Graves, S., Green, W.A., Gregor, T., Gross, N., Guerin, G.R., Günther, A., Gutiérrez, A.G., Haddock, L., Haines, A., Hall, J., Hambuckers, A., Han, W., Harrison, S.P., Hattingh, W., Hawes, J.E., He, T., He, P., Heberling, J.M., Helm, A., Hempel, S., Hentschel, J., Hérault, B., Hereş, A.M., Herz, K., Heuertz, M., Hickler, T., Hietz, P., Higuchi, P., Hipp, A.L., Hirons, A., Hock, M., Hogan, J.A., Holl, K., Honnay, O., Hornstein, D., Hou, E., Hough-Snee, N., Hovstad, K.A., Ichie, T., Igić, B., Illa, E., Isaac, M., Ishihara, M., Ivanov, L., Ivanova, L., Iversen, C.M., Izquierdo, J., Jackson, R.B., Jackson, B., Jactel, H., Jagodzinski, A.M., Jandt, U., Jansen, S., Jenkins, T., Jentsch, A., Jespersen, J.R.P., Jiang, G.F., Johansen, J.L., Johnson, D., Jokela, E.J., Joly, C.A., Jordan, G.J., Joseph, G.S., Junaedi, D., Junker, R.R., Justes, E., Kabzems, R., Kane, J., Kaplan, Z., Kattenborn, T., Kavelenova, L., Kearsley, E., Kempel, A., Kenzo, T., Kerkhoff, A., Khalil, M.I., Kinlock, N.L., Kissling, W.D., Kitajima, K., Kitzberger, T., Kjøller, R., Klein, T., Kleyer, M., Klimešová, J., Klipel, J., Kloeppel, B., Klotz, S., Knops, J.M.H., Kohyama, T., Koike, F., Kollmann, J., Komac, B., Komatsu, K., König, C., Kraft, N.J.B., Kramer, K., Kreft, H., Kühn, I., Kumarathunge, D., Kuppler, J., Kurokawa, H., Kurosawa, Y., Kuyah, S., Laclau, J.P., Lafleur, B., Lallai, E., Lamb, E., Lamprecht, A., Larkin, D.J., Laughlin, D., Le Bagousse-Pinguet, Y., le Maire, G., le Roux, P.C., le Roux, E., Lee, T., Lens, F., Lewis, S.L., Lhotsky, B., Li, Y., Li, X., Lichstein, J.W., Liebergesell, M., Lim, J.Y., Lin, Y.S., Linares, J.C., Liu, C., Liu, D., Liu, U., Livingstone, S., Llusià, J., Lohbeck, M., López-García, Á., Lopez-Gonzalez, G., Lososová, Z., Louault, F., Lukács, B.A., Lukeš, P., Luo, Y., Lussu, M., Ma, S., Maciel Rabelo Pereira, C., Mack, M., Maire, V., Mäkelä, A., Mäkinen, H., Malhado, A.C.M., Mallik, A., Manning, P., Manzoni, S., Marchetti, Z., Marchino, L., Marcilio-Silva, V., Marcon, E., Marignani, M., Markesteijn, L., Martin, A., Martínez-Garza, C., Martínez-Vilalta, J., Mašková, T., Mason, K., Mason, N., Massad, T.J., Masse, J., Mayrose, I., McCarthy, J., McCormack, M.L., McCulloh, K., McFadden, I.R., McGill, B.J., McPartland, M.Y., Medeiros, J.S., Medlyn, B., Meerts, P., Mehrabi, Z., Meir, P., Melo, F.P.L., Mencuccini, M., Meredieu, C., Messier, J., Mészáros, I., Metsaranta, J., Michaletz, S.T., Michelaki, C., Migalina, S., Milla, R., Miller, J.E.D., Minden, V., Ming, R., Mokany, K., Moles, A.T., Molnár V, A., Molofsky, J., Molz, M., Montgomery, R.A., Monty, A., Moravcová, L., Moreno-Martínez, A., Moretti, M., Mori, A.S., Mori, S., Morris, D., Morrison, J., Mucina, L., Mueller, S., Muir, C.D., Müller, S.C., Munoz, F., Myers-Smith, I.H., Myster, R.W., Nagano, M., Naidu, S., Narayanan, A., Natesan, B., Negoita, L., Nelson, A.S., Neuschulz, E.L., Ni, J., Niedrist, G., Nieto, J., Niinemets, Ü., Nolan, R., Nottebrock, H., Nouvellon, Y., Novakovskiy, A., Network, T.N., Nystuen, K.O., O’Grady, A., O’Hara, K., O’Reilly-Nugent, A., Oakley, S., Oberhuber, W., Ohtsuka, T., Oliveira, R., Öllerer, K., Olson, M.E., Onipchenko, V., Onoda, Y., Onstein, R.E., Ordonez, J.C., Osada, N., Ostonen, I., Ottaviani, G., Otto, S., Overbeck, G.E., Ozinga, W.A., Pahl, A.T., Paine, C.E.T., Pakeman, R.J., Papageorgiou, A.C., Parfionova, E., Pärtel, M., Patacca, M., Paula, S., Paule, J., Pauli, H., Pausas, J.G., Peco, B., Penuelas, J., Perea, A., Peri, P.L., Petisco-Souza, A.C., Petraglia, A., Petritan, A.M., Phillips, O.L., Pierce, S., Pillar, V.D., Pisek, J., Pomogaybin, A., Poorter, H., Portsmuth, A., Poschlod, P., Potvin, C., Pounds, D., Powell, A.S., Power, S.A., Prinzing, A., Puglielli, G., Pyšek, P., Raevel, V., Rammig, A., Ransijn, J., Ray, C.A., Reich, P.B., Reichstein, M., Reid, D.E.B., Réjou-Méchain, M., de Dios, V.R., Ribeiro, S., Richardson, S., Riibak, K., Rillig, M.C., Riviera, F., Robert, E.M.R., Roberts, S., Robroek, B., Roddy, A., Rodrigues, A.V., Rogers, A., Rollinson, E., Rolo, V., Römermann, C., Ronzhina, D., Roscher, C., Rosell, J.A., Rosenfield, M.F., Rossi, C., Roy, D.B., Royer-Tardif, S., Rüger, N., Ruiz-Peinado, R., Rumpf, S.B., Rusch, G.M., Ryo, M., Sack, L., Saldaña, A., Salgado-Negret, B., Salguero-Gomez, R., Santa-Regina, I., Santacruz-García, A.C., Santos, J., Sardans, J., Schamp, B., Scherer-Lorenzen, M., Schleuning, M., Schmid, B., Schmidt, M., Schmitt, S., Schneider, J.V., Schowanek, S.D., Schrader, J., Schrodt, F., Schuldt, B., Schurr, F., Selaya Garvizu, G., Semchenko, M., Seymour, C., Sfair, J.C., Sharpe, J.M., Sheppard, C.S., Sheremetiev, S., Shiodera, S., Shipley, B., Shovon, T.A., Siebenkäs, A., Sierra, C., Silva, V., Silva, M., Sitzia, T., Sjöman, H., Slot, M., Smith, N.G., Sodhi, D., Soltis, P., Soltis, D., Somers, B., Sonnier, G., Sørensen, M.V., Sosinski Jr, E.E., Soudzilovskaia, N.A., Souza, A.F., Spasojevic, M., Sperandii, M.G., Stan, A.B., Stegen, J., Steinbauer, K., Stephan, J.G., Sterck, F., Stojanovic, D.B., Strydom, T., Suarez, M.L., Svenning, J.C., Svitková, I., Svitok, M., Svoboda, M., Swaine, E., Swenson, N., Tabarelli, M., Takagi, K., Tappeiner, U., Tarifa, R., Tauugourdeau, S., Tavsanoglu, C., te Beest, M., Tedersoo, L., Thiffault, N., Thom, D., Thomas, E., Thompson, K., Thornton, P.E., Thuiller, W., Tichý, L., Tissue, D., Tjoelker, M.G., Tng, D.Y.P., Tobias, J., Török, P., Tarin, T., Torres-Ruiz, J., Tóthmérész, B., Treurnicht, M., Trivellone, V., Trolliet, F., Trotsiuk, V., Tsakalos, J.L., Tsiripidis, I., Tysklind, N., Umehara, T., Usoltsev, V., Vadeboncoeur, M., Vaezi, J., Valladares, F., Vamosi, J., van Bodegom, P.M., van Breugel, M., Van Cleemput, E., van de Weg, M., van der Merwe, S., van der Plas, F., van der Sande, M.T., van Kleunen, M., Van Meerbeek, K., Vanderwel, M., Vanselow, K.A., Vårhammar, A., Varone, L., Vasquez Valderrama, M.Y., Vassilev, K., Vellend, M., Veneklaas, E.J., Verbeeck, H., Verheyen, K., Vibrans, A., Vieira, I., Villacís, J., Violle, C., Vivek, P., Wagner, K., Waldram, M., Waldron, A., Walker, A.P., Waller, M., Walther, G., Wang, H., Wang, F., Wang, W., Watkins, H., Watkins, J., Weber, U., Weedon, J.T., Wei, L., Weigelt, P., Weiher, E., Wells, A.W., Wellstein, C., Wenk, E., Westoby, M., Westwood, A., White, P.J., Whitten, M., Williams, M., Winkler, D.E., Winter, K., Womack, C., Wright, I.J., Wright, S.J., Wright, J., Pinho, B.X., Ximenes, F., Yamada, T., Yamaji, K., Yanai, R., Yankov, N., Yguel, B., Zanini, K.J., Zanne, A.E., Zelený, D., Zhao, Y.P., Zheng, J., Zheng, J., Ziemińska, K., Zirbel, C.R., Zizka, G., Zo-Bi, I., Zotz, G. & Wirth, C. (2020) Try plant trait database –enhanced coverage and open access. Global Change Biology 26, 119–188.

Khedim, N., Cécillon, L., Poulenard, J., Barré, P., Baudin, F., Marta, S., Rabatel, A., Dentant, C., Cauvy-Fraunié, S., Anthelme, F., Gielly, L., Ambrosini, R., Franzetti, A., Azzoni, R.S., Caccianiga, M.S., Compostella, C., Clague, J., Tielidze, L., Messager, E., Choler, P. & Ficetola, G.F. (2021) Topsoil organic matter build-up in glacier forelands around the world. Global Change Biology 27, 1662–1677.

Kuznetsova, A., Brockhoff, P.B. & Christensen, R.H.B. (2017) lmertest package: Tests in linear mixed effects models. Journal of Statistical Software 82, 1–26.

Laliberté, E. & Legendre, P. (2010) A distance-based framework for measuring functional diversity from multiple traits. Ecology 91, 299–305.

Losapio, G., Cerabolini, B.E.L., Maffioletti, C., Tampucci, D., Gobbi, M. & Caccianiga, M. (2021) The consequences of glacier retreat are uneven between plant species. Frontiers in Ecology and Evolution 8, 1–11.

Losapio, G., de la Cruz, M., Escudero, A., Schmid, B. & Schöb, C. (2018) The assembly of a plant network in alpine vegetation. Journal of Vegetation Science 29, 999–1006.

Losapio, G. & Schöb, C. (2017) Resistance of plant–plant networks to biodiversity loss and secondary extinctions following simulated environmental changes. Functional Ecology 31, 1145–1152.

Mason, N.W., de Bello, F., Mouillot, D., Pavoine, S. & Dray, S. (2013) A guide for using functional diversity indices to reveal changes in assembly processes along ecological gradients. Journal of Vegetation Science 24, 794–806.

Mason, N.W.H., Richardson, S.J., Peltzer, D.A., de Bello, F., Wardle, D.A. & Allen, R.B. (2012) Changes in coexistence mechanisms along a long-term soil chronosequence revealed by functional trait diversity. Journal of Ecology 100, 678–689.

Mouchet, M.A., Villéger, S., Mason, N.W.H. & Mouillot, D. (2010) Functional diversity measures: An overview of their redundancy and their ability to discriminate community assembly rules. Functional Ecology 24, 867–876.

Pavoine, S. (2020) adiv: An r package to analyse biodiversity in ecology. Methods in Ecology and Evolution 11, 1106–1112.

Pérez-Harguindeguy, N., Diaz, S., Garnier, E., Lavorel, S., Poorter, H., Jaureguiberry, P., Bret-Harte, M.S., Cornwell, W.K., Craine, J.M., Gurvich, D.E., Urcelay, C., Veneklaas, E.J., Reich, P.B., Poorter, L., Wright, I.J., Ray, P., Enrico, L., Pausas, J.G., de Vos, A.C., Buchmann, N., Funes, G., Quetier, F., Hodgson, J., Thompson, K., Morgan, H.D., ter Steege, H., van der Heijden, M.G.A., Sack, L., Blonder, B., Poschold, P., Vairetti, M., Conti, G., Staver, A., Aquino, S. & Cornelissen, J.H.C. (2013) New Handbook for standardized measurment of plant functional traits worldwide. Australian Journal of Botany 61, 167–234.

RaffI, C., Mallaun, M., Mayer, R. & Erschbamer, B. (2006) Vegetation Succession Pattern and Diversity Changes in a Glacier Valley, Central Alps, Austria. Arctic, Antarctic, and Alpine Research 38, 421–428.

Reich, P.B., Wright, I.J., Cavender-Bares, J., Craine, J.M., Oleksyn, J., Westoby, M. & Walters, M.B. (2003) The evolution of plant functional variation: Traits, spectra, and strategies. International Journal of Plant Sciences 164, S143–S164.

Ricotta, C., de Bello, F., Moretti, M., Caccianiga, M., Cerabolini, B.E. & Pavoine, S. (2016) Measuring the functional redundancy of biological communities: a quantitative guide. Methods in Ecology and Evolution 7, 1386–1395.

Roches, S.D., Post, D.M., Turley, N.E., Bailey, J.K., Hendry, A.P., Kinnison, M.T., Schweitzer, J.A. & Palkovacs, E.P. (2017) The ecological importance of intraspecific variation. Nature Ecology Evolution 2, 57–64.

Rodman, K.C., Veblen, T.T., Andrus, R.A., Enright, N.J., Fontaine, J.B., Gonzalez, A.D., Redmond, M.D. & Wion, A.P. (2021) A trait-based approach to assessing resistance and resilience to wildfire in two iconic north american conifers. Journal of Ecology 109, 313–326.

Roe, G.H., Baker, M.B. & Herla, F. (2017) Centennial glacier retreat as categorical evidence of regional climate change. Nature Geoscience 10, 95–99.

Schleuning, M., García, D. & Tobias, J.A. (2023) Animal functional traits: Towards a trait-based ecology for whole ecosystems. Functional Ecology 37, 4–12.

Schleuter, D., Daufresne, M., Massol, F. & Argillier, C. (2010) A user’s guide to functional diversity indices. Ecological Monographs 80, 469–484.

Siefert, A., Violle, C., Chalmandrier, L., Albert, C.H., Taudiere, A., Fajardo, A., Aarssen, L.W., Baraloto, C., Carlucci, M.B., Cianciaruso, M.V., de L. Dantas, V., de Bello, F., Duarte, L.D.S., Fonseca, C.R., Freschet, G.T., Gaucherand, S., Gross, N., Hikosaka, K., Jackson, B., Jung, V., Kamiyama, C., Katabuchi, M., Kembel, S.W., Kichenin, E., Kraft, N.J.B., Lagerström, A., Bagousse-Pinguet, Y.L., Li, Y., Mason, N., Messier, J., Nakashizuka, T., Overton, J.M., Peltzer, D.A., Pérez-Ramos, I.M., Pillar, V.D., Prentice, H.C., Richardson, S., Sasaki, T., Schamp, B.S., Schöb, C., Shipley, B., Sundqvist, M., Sykes, M.T., Vandewalle, M. & Wardle, D.A. (2015) A global meta-analysis of the relative extent of intraspecific trait variation in plant communities. Ecology Letters 18, 1406–1419.

Song, Y., Wang, P., Li, G. & Zhou, D. (2014) Relationships between functional diversity and ecosystem functioning: A review. Acta Ecologica Sinica 34, 85–91.

Stibal, M., Bradley, J.A., Edwards, A., Hotaling, S., Zawierucha, K., Rosvold, J., Lutz, S., Cameron, K.A., Mikucki, J.A., Kohler, T.J., Šabacká, M. & Anesio, A.M. (2020) Glacial ecosystems are essential to understanding biodiversity responses to glacier retreat. Nature Ecology & Evolution 4, 686–687.

Teixeira, J.C. & Huber, C.D. (2021) The inflated significance of neutral genetic diversity in conservation genetics. Proceedings of the National Academy of Sciences 118.

Tilman, D. (1997) Mechanisms of plant competition. Plant Ecology (ed. M.J. Crawley), pp. 239–261, Blackwell Publishing Ltd.

Violle, C., Navas, M.L., Vile, D., Kazakou, E., Fortunel, C., Hummel, I. & Garnier, E. (2007) Let the concept of trait be functional! Oikos 116, 882–892.

Westerband, A.C., Funk, J.L. & Barton, K.E. (2021) Intraspecific trait variation in plants: a renewed focus on its role in ecological processes. Annals of Botany 127, 397–410.

Wright, I.J., Reich, P.B., Westoby, M., Ackerly, D.D., Baruch, Z., Bongers, F., Cavender-Bares, J., Chapin, T., Cornelissen, J.H.C., Diemer, M., Flexas, J., Garnier, E., Groom, P.K., Gulias, J., Hikosaka, K., Lamont, B.B., Lee, T., Lee, W., Lusk, C., Midgley, J.J., Navas, M.L., Niinemets, Ü., Oleksyn, J., Osada, N., Poorter, H., Poot, P., Prior, L., Pyankov, V.I., Roumet, C., Thomas, S.C., Tjoelker, M.G., Veneklaas, E.J. & Villar, R. (2004) The worldwide leaf economics spectrum. Nature 428, 821–827.

Zanzottera, M., Fratte, M.D., Caccianiga, M., Pierce, S. & Cerabolini, B.E.L. (2020) Community-level variation in plant functional traits and ecological strategies shapes habitat structure along succession gradients in alpine environment. Community Ecology 21, 55–65.

Zekollari, H., Huss, M. & Farinotti, D. (2019) Modelling the future evolution of glaciers in the european alps under the eurocordex rcm ensemble. The Cryosphere 13, 1125–1146.

Zemp, M., Frey, H., Gärtner-Roer, I., Nussbaumer, S.U., Hoelzle, M., Paul, F., Haeberli, W., Denzinger, F., Ahlstrøm, A.P., Anderson, B., Bajracharya, S., Baroni, C., Braun, L.N., Cáceres, B.E., Casassa, G., Cobos, G., Dávila, L.R., Delgado Granados, H., Demuth, M.N., Espizua, L., Fischer, A., Fujita, K., Gadek, B., Ghazanfar, A., Ove Hagen, J., Holmlund, P., Karimi, N., Li, Z., Pelto, M., Pitte, P., Popovnin, V.V., Portocarrero, C.A., Prinz, R., Sangewar, C.V., Severskiy, I., Sigurđsson, O., Soruco, A., Usubaliev, R. & Vincent, C. (2015) Historically unprecedented global glacier decline in the early 21st century. Journal of Glaciology 61, 745–762.

Zemp, M., Huss, M., Thibert, E., Eckert, N., McNabb, R., Huber, J., Barandun, M., Machguth, H., Nussbaumer, S.U., Gärtner-Roer, I., Thomson, L., Paul, F., Maussion, F., Kutuzov, S. & Cogley, J.G. (2019) Global glacier mass changes and their contributions to sea-level rise from 1961 to 2016. Nature 568, 382–386.

Zhang, J., He, N., Liu, C., Xu, L., Chen, Z., Li, Y., Wang, R., Yu, G., Sun, W., Xiao, C., Chen, H.Y.H. & Reich, P.B. (2020) Variation and evolution of c:n ratio among different organs enable plants to adapt to n-limited environments. Global Change Biology 26, 2534–2543.

