## Supplementary Information for "Glacier extinction homogenizes functional diversity"

**Table 1** Indices of parameters from mixed model testing the effects of glacier retreat (linear = '$\beta_l$', quadratic = '$\beta_q$'), species richness ('$\beta_s$'), and their interactions ('$\beta_{l,s}$','$\beta_{q,s}$') on divergence of trait means. Intercept is also reported ('$\beta_0$').

| Par | Coef | SE | CI.l | CI.h | t | $df\epsilon$ | $p$ | $r$ | $r_{CI.l}$ | $r_{CI.h}$ |
|---|---|---|---|---|---|---|---|---|---|---|
| $\beta_0$ | 0.11 | 0.01 | 0.08 | 0.15 | 9.24 | 4.62 | <0.001 | 0.97 | 0.83 | 0.99 |
| $\beta_l$ | 0.16 | 0.06 | 0.04 | 0.28 | 2.69 | 140.36 | 0.008 | 0.22 | 0.06 | 0.37 |
| $\beta_q$ | -0.19 | 0.04 | -0.28 | -0.10 | -4.39 | 141.04 | <0.001 | -0.35 | -0.47 | -0.19 |
| $\beta_s$ | ≈0.00 | ≈0.00 | ≈0.00 | ≈0.00 | 3.20 | 142.81 | 0.002 | 0.26 | 0.10 | 0.40 |
| $\beta_{l:s}$ | -0.01 | 0.00 | -0.02 | -0.01 | -3.51 | 140.47 | 0.001 | -0.28 | -0.42 | -0.13 |
| $\beta_{q:s}$ | 0.02 | 0.00 | 0.01 | 0.03 | 4.72 | 140.83 | <0.001 | 0.37 | 0.22 | 0.49 |

'Par' = parameters; 'Coef' = coefficient estimate; 'SE' = standard error of the coefficient; 'CI.l' and 'CI.h' = 95% confidence interval (lower and higher range estimate, respectively); 't' = t-value; '$df\epsilon$' = degrees of freedom of error; '$p$' = $p$-value; '$r$' = effect size; '$r_{CI.l}$' and '$r_{CI.lh}$ = 95% confidence interval of the effect size.

**Table 2** Indices of parameters from mixed model testing the effects of glacier retreat (linear = '$\beta_l$', quadratic = '$\beta_q$'), species richness ('$\beta_s$'), and their interactions ('$\beta_{l,s}$','$\beta_{q,s}$') on divergence of trait variation. Intercept is also reported ('$\beta_0$').

| Par | Coef | SE | CI.l | CI.h | t | $df\epsilon$ | $p$ | $r$ | $r_{CI.l}$ | $r_{CI.h}$ |
|---|---|---|---|---|---|---|---|---|---|---|
| $\beta_0$ | 0.16 | 0.01 | 0.14 | 0.18 | 16.75 | 8.25 | <0.001 | 0.99 | 0.95 | 0.99 |
| $\beta_l$ | 0.16 | 0.07 | 0.02 | 0.29 | 2.26 | 140.99 | 0.026 | 0.19 | 0.02 | 0.34 |
| $\beta_q$ | -0.30 | 0.05 | -0.40 | -0.20 | -5.99 | 142.49 | <0.001 | -0.45 | -0.56 | -0.31 |
| $\beta_s$ | ≈0.00 | ≈0.00 | ≈0.00 | ≈0.00 | 1.51 | 119.78 | 0.134 | 0.14 | -0.04 | 0.30 |
| $\beta_{l:s}$ | -0.01 | 0.00 | -0.02 | 0.00 | -2.43 | 141.29 | 0.017 | -0.20 | -0.35 | -0.04 |
| $\beta_{q:s}$ | 0.02 | 0.00 | 0.01 | 0.03 | 3.58 | 142.06 | <0.001 | 0.29 | 0.13 | 0.42 |

'Par' = parameters; 'Coef' = coefficient estimate; 'SE' = standard error of the coefficient; 'CI.l' and 'CI.h' = 95% confidence interval (lower and higher range estimate, respectively); 't' = t-value; '$df\epsilon$' = degrees of freedom of error; '$p$' = $p$-value; '$r$' = effect size; '$r_{CI.l}$' and '$r_{CI.lh}$ = 95% confidence interval of the effect size.

**Table 3** Indices of parameters from mixed model testing the effects of glacier retreat (linear = '$\beta_l$', quadratic = '$\beta_q$'), species richness ('$\beta_s$'), and their interactions ('$\beta_{l:s}$','$\beta_{q:s}$') on homogeneity of trait means. Intercept is also reported ('$\beta_0$').

| Par | Coef | SE | CI.l | CI.h | t | $df\epsilon$ | $p$ | $r$ | $r_{CI.l}$ | $r_{CI.h}$ |
|---|---|---|---|---|---|---|---|---|---|---|
| $\beta_0$ | 0.88 | 0.01 | 0.85 | 0.90 | 79.59 | 7.43 | <0.001 | 1.00 | 1.00 | 1.00 |
| $\beta_l$ | -0.46 | 0.09 | -0.63 | -0.29 | -5.27 | 130.32 | <0.001 | -0.42 | -0.54 | -0.27 |
| $\beta_q$ | 0.03 | 0.11 | -0.19 | 0.26 | 0.31 | 130.83 | 0.759 | 0.03 | -0.14 | 0.19 |
| $\beta_s$ | ≈0.00 | ≈0.00 | ≈0.00 | ≈0.00 | -2.93 | 123.76 | 0.004 | -0.25 | -0.40 | -0.08 |
| $\beta_{l:s}$ | 0.02 | 0.01 | 0.01 | 0.03 | 3.45 | 130.97 | 0.001 | 0.29 | 0.12 | 0.43 |
| $\beta_{q:s}$ | ≈0.00 | 0.01 | -0.02 | 0.01 | -0.39 | 131.16 | 0.699 | -0.03 | -0.20 | 0.14 |

'Par' = parameters; 'Coef' = coefficient estimate; 'SE' = standard error of the coefficient; 'CI.l' and 'CI.h' = 95% confidence interval (lower and higher range estimate, respectively); 't' = t-value; '$df\epsilon$' = degrees of freedom of error; '$p$' = $p$-value; '$r$' = effect size; '$r_{CI.l}$' and '$r_{CI.lh}$ = 95% confidence interval of the effect size.

**Table 4** Indices of parameters from mixed model testing the effects of glacier retreat (linear = '$\beta_l$', quadratic = '$\beta_q$'), species richness ('$\beta_s$'), and their interactions ('$\beta_{l,s}$','$\beta_{q,s}$') on homogeneity of trait variation. Intercept is also reported ('$\beta_0$').

| Par | Coef | SE | CI.l | CI.h | t | $df\epsilon$ | $p$ | $r$ | $r_{CI.l}$ | $r_{CI.h}$ |
|---|---|---|---|---|---|---|---|---|---|---|
| $\beta_0$ | 0.89 | 0.01 | 0.87 | 0.91 | 95.40 | 9.57 | <0.001 | 1.00 | 1.00 | 1.00 |
| $\beta_l$ | 0.18 | 0.09 | 0.01 | 0.36 | 2.11 | 130.26 | 0.036 | 0.18 | 0.01 | 0.34 |
| $\beta_q$ | 0.13 | 0.11 | -0.09 | 0.35 | 1.15 | 131.12 | 0.253 | 0.10 | -0.07 | 0.26 |
| $\beta_s$ | ≈0.00 | ≈0.00 | ≈0.00 | ≈0.00 | -1.25 | 95.47 | 0.214 | -0.13 | -0.31 | 0.07 |
| $\beta_{l:s}$ | 0.01 | 0.01 | -0.01 | 0.02 | 0.99 | 131.39 | 0.322 | 0.09 | -0.08 | 0.25 |
| $\beta_{q:s}$ | -0.02 | 0.01 | -0.03 | 0.00 | -2.35 | 131.64 | 0.020 | -0.20 | -0.35 | -0.03 |

'Par' = parameters; 'Coef' = coefficient estimate; 'SE' = standard error of the coefficient; 'CI.l' and 'CI.h' = 95% confidence interval (lower and higher range estimate, respectively); 't' = t-value; '$df\epsilon$' = degrees of freedom of error; '$p$' = $p$-value; '$r$' = effect size; '$r_{CI.l}$' and '$r_{CI.lh}$ = 95% confidence interval of the effect size.
